# The TLR5 agonist flagellin modifies phenotypical and enhances functional activation of lung mucosal antigen presenting cells in neonatal mice

**DOI:** 10.1101/564054

**Authors:** Pankaj Sharma, Ofer Levy, David J. Dowling

**Affiliations:** Precision Vaccines Program, Division of Infectious Diseases, Boston Children’s Hospital, Boston, MA, USA; Harvard Medical School, Boston, MA, USA; Broad Institute of MIT and Harvard

**Author notes:** Co-senior and Co-corresponding authors:David J. Dowling,Ofer LevyEmail. Footnote: The *Precision Vaccines Lab* is supported by U.S. National Institutes of Health (NIH)/National Institutes of Allergy and Infectious Diseases (NIAID) awards including Molecular Mechanisms of Combination Adjuvants (1U01AI124284-01), Adjuvant Discovery (HHSN272201400052C) and Development (HHSN272201800047C) Program Contracts and an internal Boston Children’s Hospital award. DD’s laboratory is supported by NIH grant 1R21AI137932-01A1.

**Keywords:** Early life immunization, newborn, dendritic cells, mucosal immunity, cross presentation, TLR5, flagellin

## Abstract

Intranasal mucosal vaccines are of interest in that they may induce protective mucosal immune responses. Activation of lung antigen presenting cells (APCs), a phenotyoically and functionally heterogeneous cell population located at distinct mucosal sites, may be key to the immunogenicity of such vaccines. Characterizing responsiveness of newborn lung APCs to adjuvants may inform design of efficacious intranasal vaccines for early life, when most infections occur. We characterized APCs from neonatal (<7 days of life) and adult (6-8 weeks of age) mice. Neonatal mice displayed a relatively high abundance of alveolar macrophages (AMs), with lower percentages of plasmacytoid dendritic cells (pDCs), CD103^+^ (cDC1) and CD11b^+^ (cDC2) DCs. Furthermore, neonatal CD103^+^ and CD11b^+^ DC subsets demonstrated an inverse expression of maturation markers as compared to adult mice. Upon stimulation of lung APC subsets with a panel of pattern recognition receptor (PRR), including TLR and STING, agonists, CD11c^+^ enriched cells from neonatal and adult mice lungs demonstrated distinct maturation profiles. The TLR5 ligand, flagellin, was most effective at activating neonatal lung APCs, inducing significantly higher expression of maturation markers on CD103^+^ (cDC1) and CD11b^+^ (cDC2) subsets. Intranasal administration of flagellin induced a distinct migration of CD103^+^ and CD11b^+^ DC subsets to the mediastinal lymph nodes (mLNs) of neonatal mice. Overall, these findings highlight age specific differences in the maturation and responsiveness of lung APC subsets to different PRR agonists. The unique efficacy of flagellin in enhancing lung APC activity suggests that it may serve as an effective adjuvant for early life mucosal vaccines.

## Introduction

The persistently high global burden of infections in the very young provides a compelling rationale for developing additional safe and effective early life vaccines (1). Most childhood pathogens access the body through mucosal membranes (2). Additionally, viral infections including respiratory syncytial virus (RSV) and influenza virus are often more severe and/or prolonged in early life as compared to adult life (3). Parenterally vaccines, such as those delivered intramuscularly, are often poor inducers of protective immunity at mucosal surfaces (4). While, effective immunization against mucosal infections usually requires topical-mucosal vaccine administration, which could neutralize the pathogen on the mucosal surface before it can cause infection, only a few human vaccines are oral (against cholera, typhoid, polio, and rotavirus), and only one is administered intranasal (against influenza) (5). Moreover, vaccines delivered via different routes interact with different APCs that activate different arms of immunity (6). Tissue distribution and migratory properties of APCs, such as alveolar macrophages (AMs), plasmacytoid DCs (pDCs), and conventional DCs (cDCs), contribute to the generation of distinct immune responses (7). In mice, cDCs have been classified into two major subsets, cDC1, which express CD103 (CD8_α_) and specialize in cross-presentation to CD8^+^ T cells critical for immunity against intracellular pathogens, viruses, and cancer; and cDC2, which express CD11b and promote CD4^+^ T cell differentiation into subsets specializing in anti-viral, - fungal, or -helminth immunity (8). Immune cells in the neonatal lung differ in quantity and quality from adults and, hence, react differently to environmental, microbial and vaccine exposures (9). Growing evidence suggests that mucosal administration of antigen may drive a more effective mucosal response to respiratory infections (10,11). This may be partly due to the activation of antigen-specific secretory IgA responses and development of lung resident memory T cells (12-14).

Generation of mucosal immunity can potentially be achieved via various routes including oral, intranasal (pulmonary), rectal, and vaginal (4). The intranasal route is attractive considering injection-free delivery (15), and ease of clinical administration (16) in early life. However, the challenge to early life intranasal vaccination can be the induction of nonspecific inflammation and/or the generation of tolerance to unknown or novel antigens (17). Such challenges may be a particular concern in an early life, where the T-helper response is biased toward type-2 immunity, exacerbation of which may predispose individuals to eosinophilia and pulmonary disorders (18). Nevertheless, solutions to such challenges may be found in the use of particular adjuvants which drive the appropriate activation of APCs and help in shaping the immune system (19,20). Adjuvantation is a key tool to enhance vaccine-induced immunity (21). Adjuvants can enhance, prolong, and modulate immune responses to vaccine antigens to maximize protective immunity (20,21), and may potentially enable more effective immunization in the very young and the elderly (22). Despite some evidence suggesting ontological differences in the innate response in the mucosal APCs, there are gaps in the understanding of basic mechanisms and if/which known adjuvants could drive mature and enhance the functions of lung APC subsets in diverse age groups such as infant and neonates (23). A deeper understanding of mucosal APCs in early life may inform design of effective mucosal vaccines for the very young.

In the present study, we examined the phenotypic and functional differences in the mucosal APC subsets isolated from newborn and adult murine lungs and characterized their responsiveness to different pattern recognition receptor (PRR) agonists/adjuvants. We found that DC subsets from neonate mice lungs are phenotypically and functionally distinct from those of the adult mice, as they exhibit a different activation pattern to various pattern recognition stimuli. The TLR5 agonist flagellin, a globular protein that arranges itself in a hollow cylinder to form the filament in a bacterial flagellum, strongly activated lung migratory DC (migDCs) subsets and upregulated their expression of CD40, CD80, CD86 and CCR7. Also, when used as an adjuvant during intranasal vaccination, flagellin regulated the migration of DC subsets to the draining lymph nodes, and potentiated the phagosome maturation in CD103^+^ DC subset, correlating with their ability to mount a robust antigen cross presentation phenontype. These findings suggest that TLR5 may play an important role in the maturation and activation of lung migratory DCs in neonates and could be a empirically promising target for intranasal mucosal vaccines delivered in early life.

## Methods

### Animals

All experiments involving animals were approved by the Animal Care and Use Committee of Boston Children’s Hospital and Harvard Medical School (protocol numbers 15-11-3011 and 16-02-3130). C57BL/6 mice were obtained from Taconic Biosciences or Charles River Laboratories and housed in specific pathogen-free conditions in the animal research facilities at Boston Children’s Hospital. For breeding purposes, mice were housed in couples, and cages checked daily to assess pregnancy status of dams and/or the presence of pups. When a new litter was discovered, that day was recorded as day of life (DOL) 0. Both male and female pups were used for experiments.

### Reagents

Collagenase-1, DNase-1, MTT, Griess reagent, FITC-dextran, and Concanavalin-A were all purchased from Sigma Aldrich (St Louis, MO). CD11c magnetic cell separation kit (MACS) were purchased from Miltenyi Biotec (Bergisch-Gladbach, Germany). All monoclonal antibodies (Abs) were purchased from eBioscience (San Diego, CA), while all the PRR agonists were purchased from Invivogen (San Diego, CA). For all *in vivo* studies, ultrapure Salmonella typhimurium flagellin (FLA-ST) from invivogen (endotoxin level < 0.05 EU/mg) and was employed. FLA-ST is purified by acid hydrolysis, heating and ultrafiltration to obtain an estimated at 10% purity, before an additional purification step using monoclonal anti-flagellin affinity chromatography to obtain purity > 95%.

### Preparation of single cell suspension from murine lungs and isolation of lung APCs

To prepare single cell suspensions, neonates and adult mice were sacrificed and lungs were exposed by dissecting through the thorax by raising the sternum to avoid any injury to the lung. Mice lungs were perfused through right ventricle by adding ice cold PBS (containing 2 mM EDTA) and lung lobes were carefully isolated and were manually minced into small pieces in petri dishes. Next, these minced pieces were incubated at 37°C for 30 min in a digestion medium containing 2mg/ml of collagenase and 80U/ml of DNase-1. Digested lung tissue was subsequently passed through a 40 µM cell strainer and centrifuged at 300 g for 10 min at 4°C. RBC lysis was carried out in the cell pellet using BD Pharm Lyse following instructions of the manufacturer (BD Biosciences). For multicolor immunophenotyping and in vitro stimulation assay, lung APCs were enriched using CD11c based magnetic activated cell sorting (MACS) following the manufacturer’s instructions.

### *In vitro* stimulation assay

All PRR agonists employed in the studies were verified endotoxin-free as indicated by the manufacturers (Invivogen). For stimulation experiments, isolated lung CD11c^+^ cells from newborn and adult mice were plated in round bottom 96-wells non-tissue culture-treated plates at the density of 10^5^ cells/well in 200 µl of fresh complete culture medium and were stimulated with 100 ng/ml of PAM3CSK4, PAM2CSK4, Poly I:C, MPLA, S. typhimurium Flagellin, 5’ppp-dsRNA, or 0.1 μM of CL075, CpG class C - ODN 2395 100 μg/ml of 2’3’-cGAMP as previously described (24,25). All Abs used for flow cytometry are listed in Table S1.

### Flow cytometry

CD11c enriched cells from mice lungs were stained with the monoclonal Abs directed against CD11c, CD11b, CD103, F4/80 and PDCA-1 to identify the five major APC populations in the lung. To monitor the expression of co-stimulatory molecules, *in vitro* stimulated CD11c^+^ cells were washed with ice cold PBS once and were thereafter stained with Ab cocktail containing monoclonal Abs for CD40, CD80, and CD86 along with APC surface markers. Data were acquired on BD Fortessa flow cytometer and was analyzed using Flowjo (Treestar).

### Analysis of phagocytosis and antigen processing capacity of lung APCs

Isolated CD11c^+^ cells were incubated with either FITC**-**dextran (1 mg/ml) or with DQ™ ovalbumin (DQ-OVA) (0.5 mg/ml) for 45 min in a CO2 incubator at 37^°^C and the fluorescence of gated cells was measured using BD Fortessa as described recently (26).

### Intranasal administration of DQ-OVA

50 µl of PBS, DQ-OVA (50 µg), and flagellin (3 μg total) adjuvanted DQ-OVA (50 µg) were applied *via* the nostrils as recently described (27). DQ-OVA was used for degradation and accumulation assays as it consists of OVA bound to a self-quenching fluorescent dye, which upon intracellular degradation releases specific fluorescence (excitation at 505□nm, emission at 515□nm). Accumulated DQ-OVA forming dimers emit fluorescence in a different channel (excitation at 488□nm, emission at 613□nm). Animals were euthanized 4 or 24□h after intranasal administration and lung and or lung draining lymph nodes were harvested to determine the uptake, trafficking, phenotype and antigen degradation (27). To enumerate the antigen cross presentation, DCs from the draining lymph nodes were stained with anti-mouse Kb-SIINFEKL Abs, and data was acquired on BD Fortessa flow cytometer.

### Statistical analysis

Statistical significance and graphs were generated using Prism v. 7.0a (GraphPad Software, La Jolla, CA, USA) and Microsoft Excel (Microsoft Corporation, Redmond, WA). For data analyzed by normalization to control values (vehicle), column statistics were conducted using the two-tailed Wilcoxon Signed Rank Test or unpaired Mann-Whitney test as appropriate. Group comparisons employed two-way ANOVA with Sidak multiple comparison post-test. Results were considered significant at *p* < 0.05, and indicated as follows: \**p*<0.05, \*\**p*<0.01, \*\*\**p*<0.001, \*\*\*\**p*<0.001.

## Results

### Neonate murine lung antigen presenting cells and their precursors demonstrate distinct percentages and co-stimulatory molecule expression

To prepare single cell suspensions, neonates and adult mice were sacrificed and lungs isolated, digested and enriched for CD11c^+^ cell fractions (Fig. 1A). A panel of reliable and cell surface makers were used in combination to identify heterogeneous populations of lung APCs studied in this article (Fig. 1B-C). As mucosal APCs are heterogeneous, we employed a stringent gating strategy to distinguish the different subsets and their progenitors. By using the combination of different surface markers and a hierarchal gating strategy, we identified alveolar macrophages (AMs), moDCs, CD103^+^ DCs, CD11b^+^ DCs and pDCs (Fig. 2A). While the percentage of Ly6C^+^ cells was significantly higher in neonatal vs. adult mice (Fig. 2B), and AMs and moDCs subsets were similar between both age groups, neonates demonstrated a significanly lower percentage of lung DC subsets CD103^+^ DCs (n = 12, *P* < 0.001), CD11b^+^ DCs (*P* < 0.001) and pDCs (*P* < 0.05) (Fig. 2C). Quite interestingly, while both adult and neonatal antigen uptake capability was similar for all the major APC subsets (Figure 3A), CD103^+^ and CD11b^+^ DCs from neonates showed relatively lower expression of basal co-stimulatory molecules CD40 (CD11b^+^ (n = 12, *P* < 0.001) and CD103^+^ (*P* < 0.01) DCs), CD80 (*P* < 0.001) and CD86 (*P* < 0.001) (Fig. 3C-D). These observations demonstrated that neonatal murine lung APCs contain a greater proportion of phenotypically distinct immature DC precursor cells, including CD103^+^ and CD11b^+^ DCs, relative to their adult counterparts.

**Figure 1.**
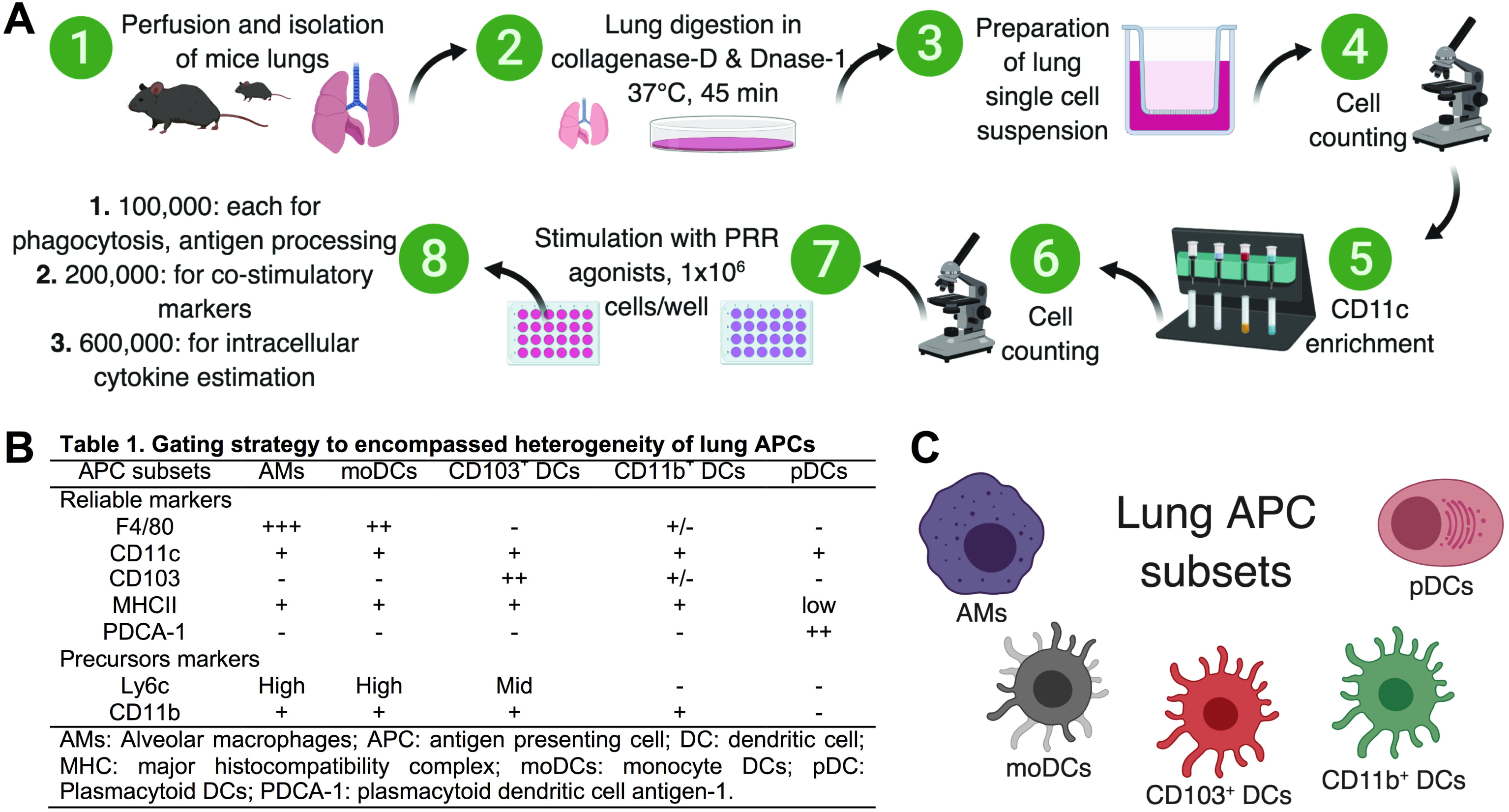
Overview of the preparation of single cell suspension from neonatal and adult murine lungs and isolation of lung APCs. (**A**) To prepare single cell suspensions, neonates and adult mice were sacrificed and lungs were exposed by dissecting through the thorax by raising the sternum to avoid any injury to the lung, perfused through right ventricle by adding ice cold PBS (containing 2mM EDTA) and lung lobes were carefully isolated and were manually minced into small pieces in petri dishes. Minced pieces were digested and digested lung tissue was subsequently passed through cell strainer. For multicolor immunophenotyping and *in vitro* stimulation assay, lung APCs were enriched using CD11c based magnetic beads for use in in vitro stimulation assays. (**B**) List of reliable and precursor makers used in combination to identify heterogeneous populations of lung APCs. (**C**) Overview of the lung APCs studied in this article.

**Figure 2.**
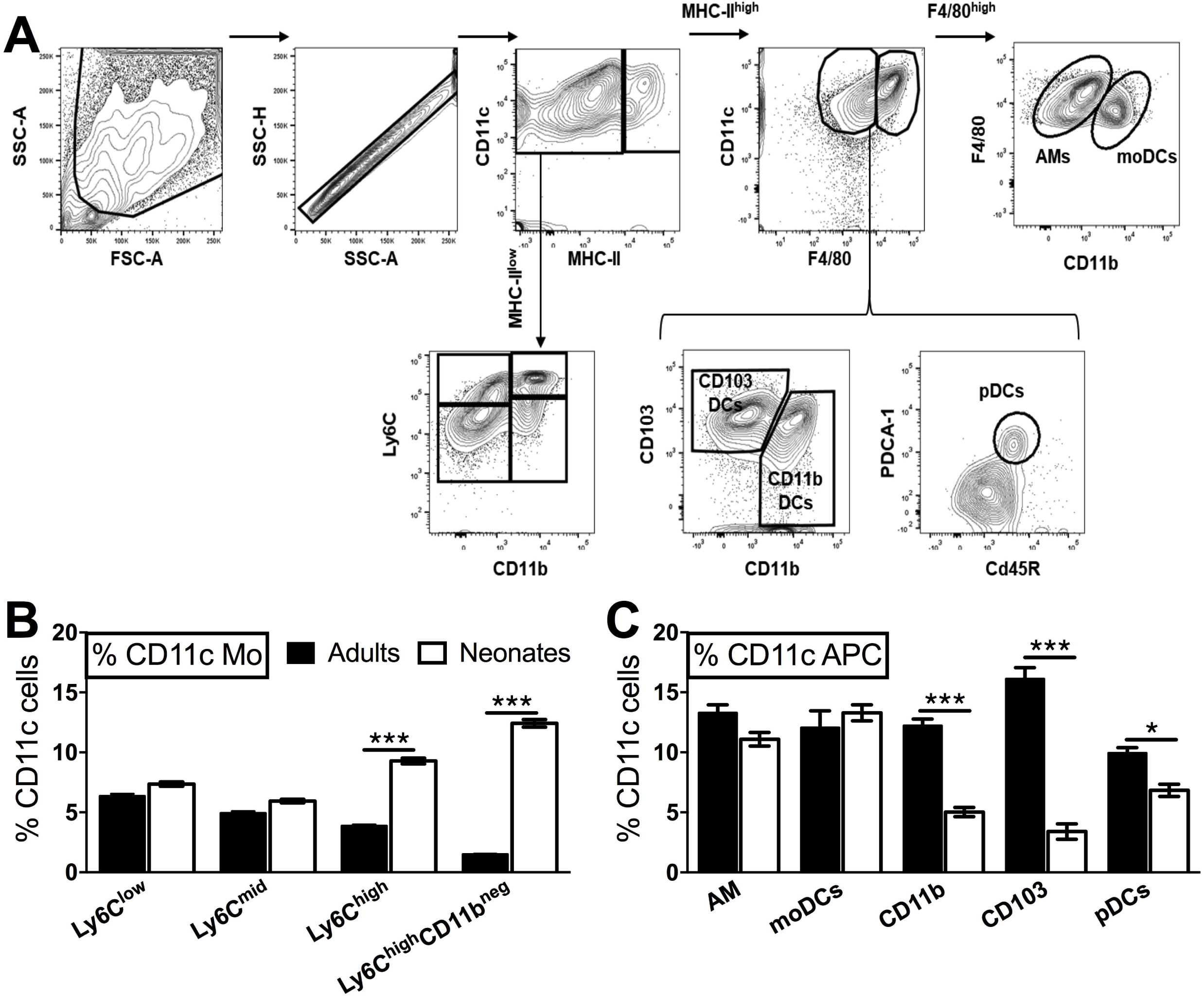
Neonate murine lung antigen presenting cells and their precursors demonstrate distinct subset percentages. (**A**) An overview of the flow cytometry gating strategy used for the identification of murine APC precursors and subsets in both adult and neonatal lungs. (**B**) The percentage of APC precursors, as defined by CD11c, Ly6C and CD11b cell expression. (**C**) The percentage of APC subsets, as defined by CD11c, F4/80, CD11b, CD103 PDCA-1 and CD45R cell expression in mice lungs. Results are expressed as mean ± SEM of n = 12 per age group. *p < 0.05, ***p < 0.001 determined by repeated measures two-way ANOVA with Sidak post hoc test.

**Figure 3.**
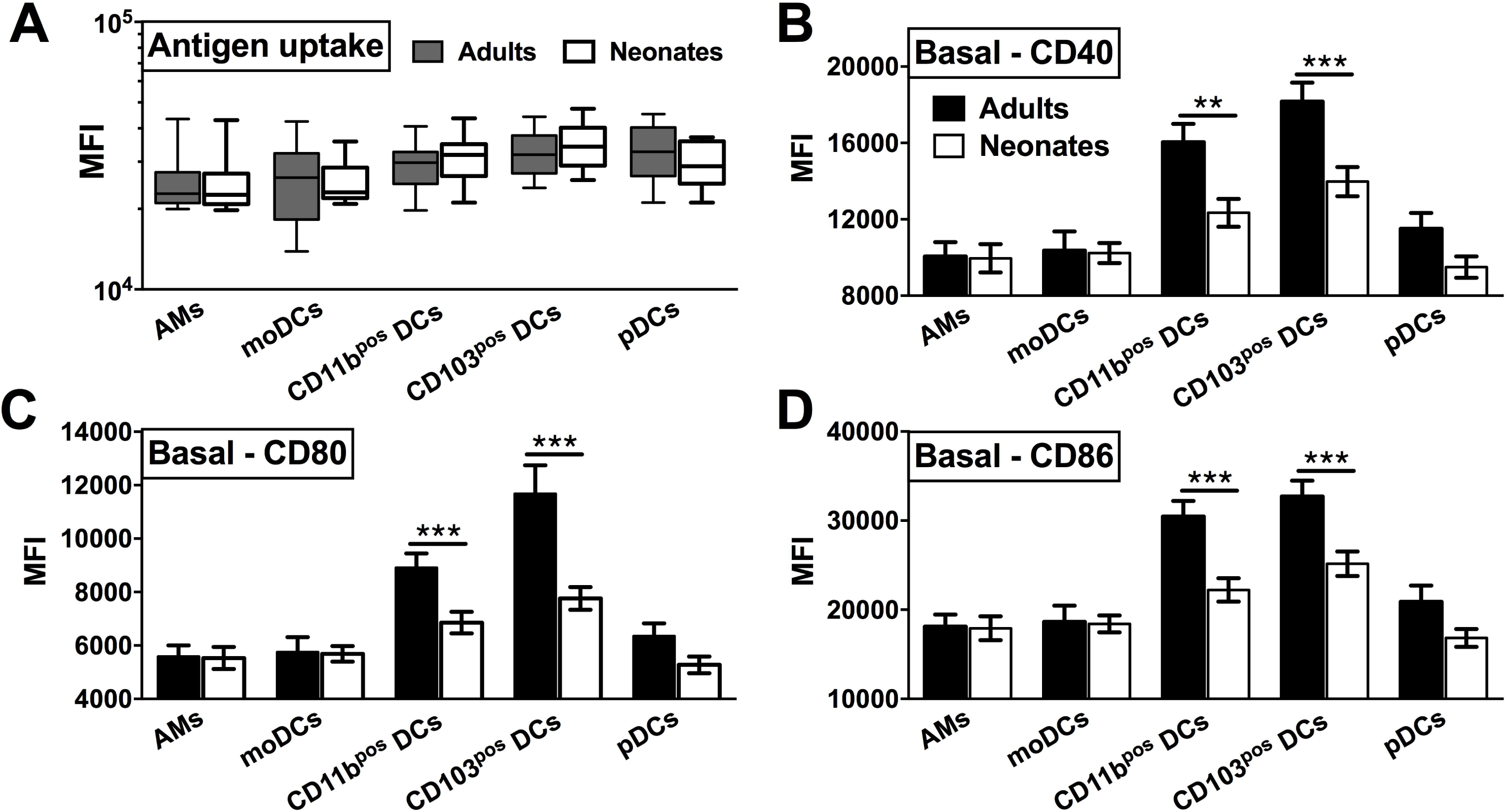
Neonatal murine lung CD103+ DCs and CD11b+ DC subsets demonstrate distinct basal co-stimualtory molecule expression. (**A**) No ontogeny based differences are observed in the antigen uptake capacity across all the DC subsets. (**B-D**) Neonatal CD103+ DCs and CD11b+ lung DC subsets demonstrate and inverse expression of co-stimulatory molecules CD40, CD80, and CD86 as compared to adult mice. Results are expressed as mean ± SEM of n = 12 per age group. *p < 0.05, ***p < 0.001 determined by repeated measures two-way ANOVA with Sidak post hoc test.).

### Neonatal murine lung APCs demonstrate distinct functional responses to PRR agonists

Our observations regarding the phenotype of DC subsets in neonatal murine lungs indicated an intrinsic age-specific difference in the percentages and maturation status of migratory APC subsets, raising the possibility that functional responses of migDCs subsets to stimuli such as PRR agonist adjuvants may also be distinct. To test this hypothesis, we stimulated CD11c^+^ cells isolated from the lungs of different aged mice and stimulated them with different classes of PRR receptor agonists at a concentrations reported most active for bone marrow DC activation (24,25). We then summarized the combined flow cytometry all APCs maturation data collected following *ex vivo* stimulation, and graphed the data as volumetric dot sizes indicating the relative fold change (ranging form < 1 to > 4.5) for CD40, CD80 and CD86, per F4/80, CD11b, CD103 PDCA-1 APC subsets, as compared with un-stimulated controls per age group. Overall, unique activation and maturation patterns were observed by stimuli, age, marker and APC subset (Fig. 4). Ranking by sum fold change activation, CpG ODN ranked first in adult mice. Flagellin ranked first in the ability to mature neonatal lung APCs (Fig. 4). While the responses of mucosal APC subsets to PRR agonists TLR1 (PAM3CSK), TLR2 (PAM2CSK), and TLR3 (Poly I:C) were impaired in general, we noted that all the neonatal APCs responded very poorly to NOD1 and NOD2 ligands (C12-iE-DAP and L18-MDP), which moderately activated the migDC subsets and pDCs from adult mice lings. Contrary to these ligands, TLR5 agonist flagellin and STING agonist 2’3’-cGAMP specifically and strongly activated the neonatal CD103^+^ and CD11b^+^ DC subsets, while the TLR4 (MPLA), TLR8/7 (CL075), TLR9 (CpG) and RIG-I (5’ppp-dsRNA) showed comparable activity for all the DC subsets from the neonates and adult mice. These observations demonstrate functional differences in the major APC subsets of murine lungs and highlight flagellin and 2’3’-cGAMP as a potential activators of mucosal immunity in early life.

**Figure 4.**
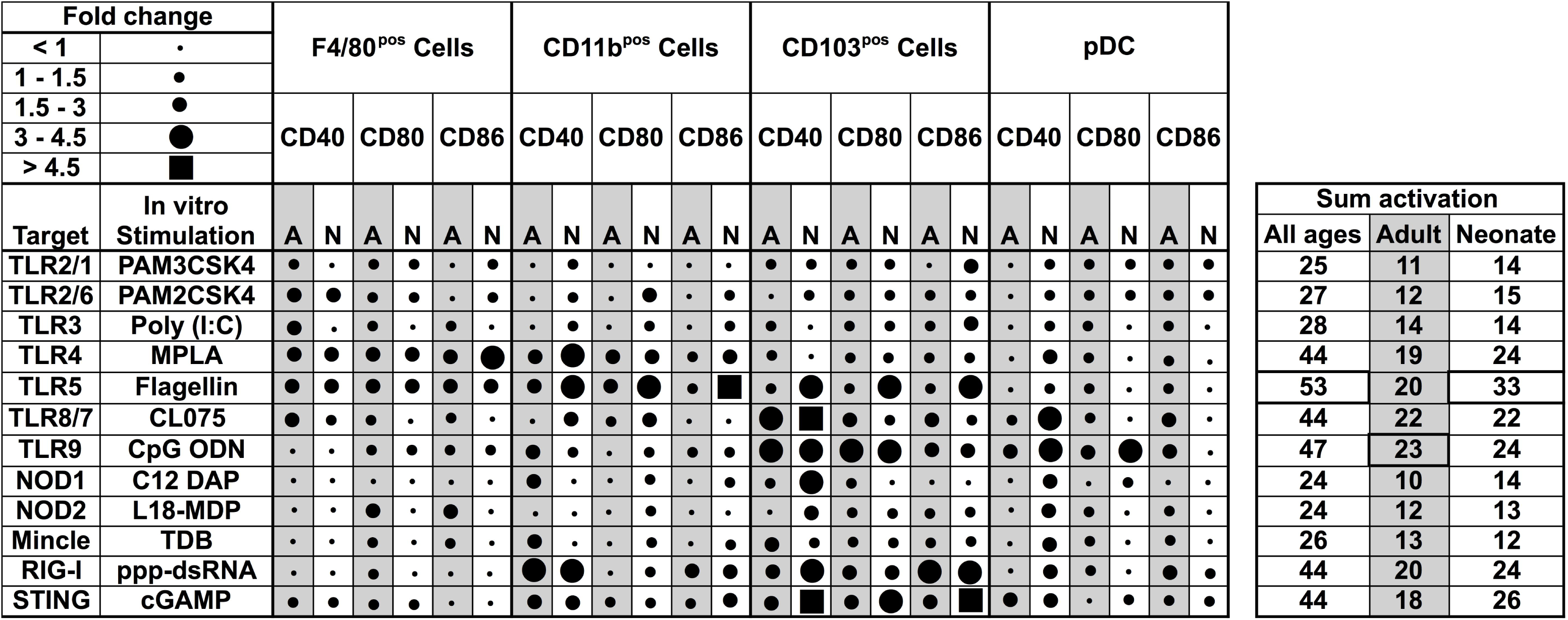
Functional differences in the major APC subsets in mice lungs identify out flagellin and 2’3’-cGAMP as a potential activators of mucosal immunity in early life. Summary of flow cytometry all APCs maturation data collected ex vivo stimulation. Dot sizes indicate the fold change (ranging form < 1 to > 4.5) for CD40, CD80 and CD86, per F4/80, CD11b, CD103 PDCA-1 APC subsets, as compared with un-stimulated controls per age group. Gray-shaded sections indicate adult data. Combined activation totals fold changes are summarized in the right panel. While the responses of mucosal APC subsets to PRR agonists TLR2/1 (PAM3CSK4), TLR2/6 (PAM2CSK4), and TLR3 (Poly I:C) were impaired in general, neonatal APCs responded very poorly to NOD1 and NOD2 ligands (C12-iE-DAP and L18-MDP respectively), which moderately activated the migDC subsets and pDCs from adult mice lings. Contrary to these ligands, TLR5 agonist flagellin and STING agonist 2’3’-cGAMP specifically and strongly activated the neonatal CD103^+^ and CD11b^+^ DC subsets, while the TLR4, TLR8/7, TLR9 and RIG-I (5’ppp-dsRNA) showed comparable activity for all the DC subsets from the neonates and adult mice. n = 5 per age group.

### Flagellin upregulates the expression of CCR7 on neonatal CD11c enriched cells and drives the lymph node homing of migDC subsets

As different stimuli induce distinct activation of lung APC subsets, we characterized the expression of lymph node homing receptor CCR7 on lung CD11c^+^ cells isolated from the neonatal and adult mice. CCR7 and CCR9 are both chemokine receptors that regulate homing of immune cells. Specifically, up-regulation of CCR7 on mature DCs is required for migration to draining/secondary lymphoid organs (28,29). Conversely, immature DCs express high levels of CCR9, as an indication of lower migration potential (30). Consistent with the *in vitro* co-stimulatory molecule stimulation data, and when ranked by MFI, MPLA, R848, CpG-ODN and STING demonstrated moderate upregulation of CCR7 on neonates as well as on the adult APCs (Fig. 5A), while as compared to unstimulated controls, flagellin most strongly upregulated the expression of CCR7 on neonatal APCs (Fig. 5B, (*P* < 0.001)) and to a significant but lesser level in adults (*P* < 0.01). Conversely, as compared to adults (*P* < 0.05), neonatal lung CD11c+ DCs CCR9 expression was not significantly increased (Fig. 5C). We also noted that the APCs from the neonates did not respond to the NOD1 and NOD2 agonists studied (C12-iE-DAP and L18-MDP) (Fig. 5A). These findings suggest that TLR5 might drive the migration of DC subsets to draining lymph nodes and initiate adaptive immunity.

**Figure 5.**
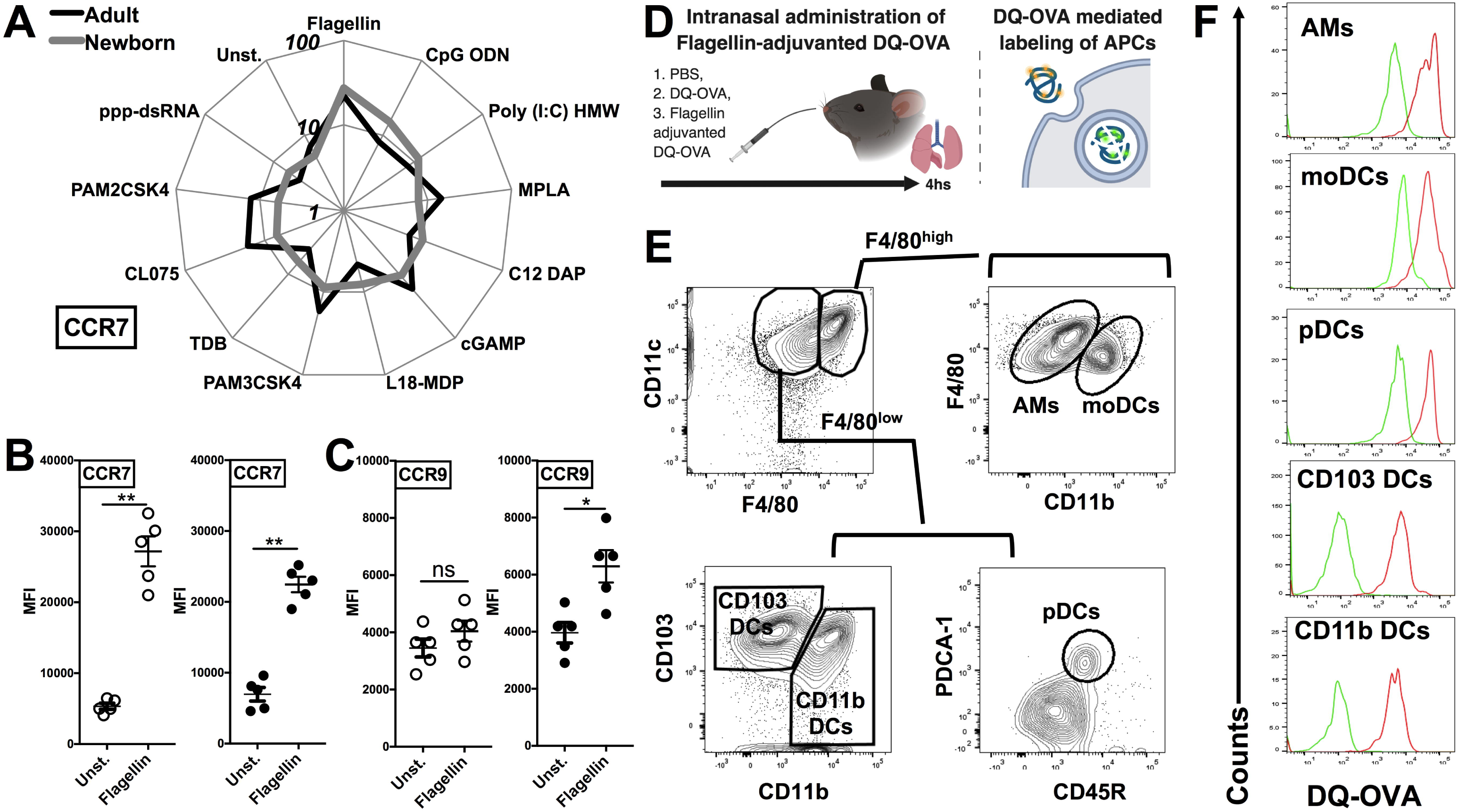
Flagellin upregulates the expression of CCR7 on neonatal CD11c enriched cells and drives the lymph node homing of migDC subsets. (**A**) Characterization of lung CD11c+ cells isolated from the neonatal and adult mice, identifies TLR5 agonist flagellin as the most robust inducer of the expression of lymph node homing receptor CCR7 (MFI ranked clockwise high (flagellin) to low (unstimulated)). (**B-C**) Flagellin induced CCR7 expression is inversely related to the expression of CCR9 on neonatal lung CD11c+ DCs. (**D**) Overview of Intranasal administration of Flagellin-adjuvanted DQ-OVA and DQ-OVA mediated labeling of APCs. DQ-OVA is a self-quenched conjugate of ovalbumin antigen that exhibits bright green fluorescence signal only upon proteolytic cleavage/degradation by endosomal proteases. (E) Gating strategy for identification of APC subsets in mice lungs. (F) DQ-OVA uptake by lung APC subsets 4 hrs after intranasal administration. n = 5 per age group. For analyses of individual treatments (e.g., unstim vs. flagellim), the unpaired Mann-Whitney test was applied at each concentration, and statistical significance denoted as *p < 0.05, **p < 0.01.

To test this hypothesis, we administrated the DQ-OVA antigen either alone or in combination with flagellin by intranasal route and measured migration of lung DCs to draining lymph nodes (Fig. 5D). DQ-OVA is a self-quenched conjugate of ovalbumin antigen that exhibits bright green fluorescence signal only upon proteolytic cleavage/degradation by endosomal proteases (31) (Fig. 5D). Since, as we outlined in Figure 2A, DC subsets from neonates and adult showed no significant differences in the antigen uptake, we could rely on DQ-OVA mediated labeling for these studies. Also, to rule out any difference which might be because of accessibility of DQ-OVA to lung interstitial resident DCs (moDCs and CD11b DCs), we looked at DQ-OVA uptake by lung APC subsets 4 hrs after intranasal administration, and our data clearly data demonstrated that all the major APC subsets were labeled (Figure. 5E-F).

Next, to examine trafficking of leukocytes, we sacrificed animals 24 hrs after intranasal administration of DQ-OVA and isolated the draining lymph nodes (Figure. 6A). In the lymph node single cell suspension, we gated lung migratory DCs as CD11c^+^, MHC-II^high^ and DQ-OVA^green+^ cells (Fig. 6B) (32). Neonates compared to adult demonstrated lower basal percentages of both the cDC subsets, while mice receiving flagellin-adjuvanted DQ-OVA demonstrated higher trafficking of immunogenic migDCs (Fig. 6C). Notably, flagellin also increased the percentage of adult CD11b^+^ cells in mLNs (Fig. 6C). Moreover, flagellin upregulated the expression of co-stimulatory molecules on neonatal CD103^+^ DCs, while no significant differences were observed on the other subset studied (Figure. 6D-E). These observations clearly show that TLR5 agonist adjuvantation mediated stimulation potentially activates the lung CD103^+^ and CD11b^+^ migDC subsets in neonates.

**Figure 6.**
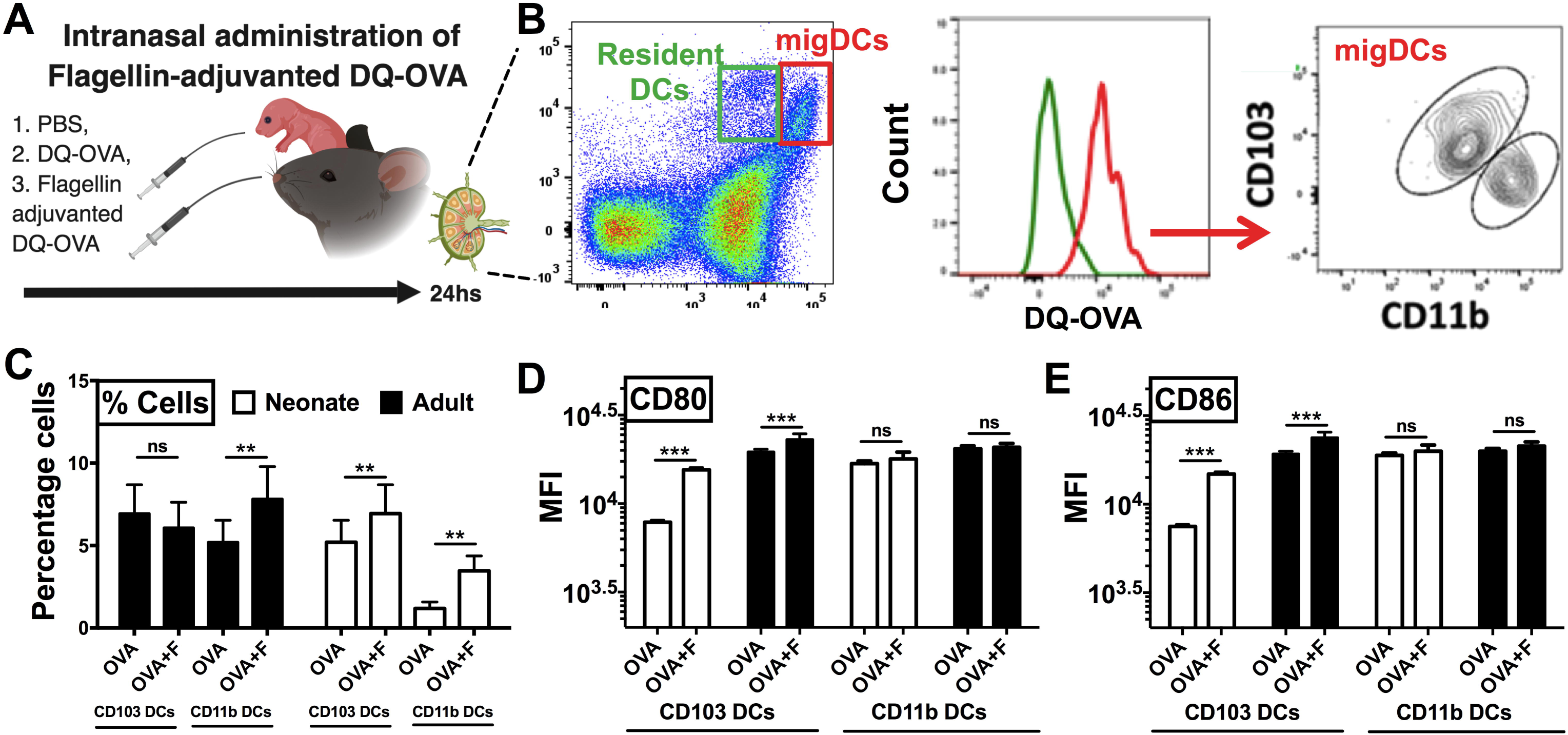
Neonatal mice exhibited a comparative polarized migration of CD103^+^ and CD11b^+^ DC subsets to the mediastinal lymph nodes (mLNs) after intranasal administration of flagellin. (**A**) Overview of Intranasal administration of Flagellin-adjuvanted DQ-OVA and isolation of draining lymph nodes at 24 hrs. (**B**) Gating strategy for identification of migratory DC1s and DC2s in draining lymph nodes. (**C**) Flagellin increases the percentage of OVA specific adult CD11b+ cells in mLNs, but both CD103+ and CD11b+ DC subsets in neonates. (**D**) Flagellin upregulates the expression of CD80 and CD86 on neonatal CD103+ migDCs DCs isolated from mLNs. Results are expressed as mean ± SEM of n = 5 per age group. *p < 0.05, ***p < 0.001, ns, not significant, determined by repeated measures two-way ANOVA with Sidak post hoc test).

### Flagellin regulates phagosome maturation and antigen degradation by neonatal lung CD103^+^ DCs

By using DQ-OVA, we not only quantified cell trafficking but also characterized antigen uptake, degradation, and cytosolic distribution. DQ-OVA gives green fluorescence in a green channel, and the co-emergence of a red signal reflects the accumulation of cleaved OVA peptides enabling assessment of cytoplasmic differences in organelle distribution after endocytosis. First, we stimulated lung CD11c^+^ cells *in vitro* with flagellin and measured the phagosome accumulation of DQ-OVA peptides (Fig. 7A). We noted the dramatic reduction in DQ-OVA^red^ signal in neonates CD103^+^ DCs in a time dependent manner, which was corrected by flagellin stimulation (Fig. 7A, (*P* < 0.001)). In line with our *in vitro* studies, we explored this process *in vivo* using our LN migration model. Once again, the flagellin adjuvanted group demonstrated increased OVA^red^ signal, but only in neonatal, and not adult (not shown), CD103^pos^ DCs (Fig. 7B, (*P* < 0.001)). Notably, we did not observe any significance downregulation or impaired phagosome maturation in either adult CD103^+^ or CD11b^+^ DCs as compared to their neonatal counterparts subsets (not shown). As slow phagosome maturation is a key step for antigen cross-presentation, these observations suggest that flagellin adjuvant in neonates, but not necessarily adult mice, might enhance a mucosal resident cytotoxic T cell response. These findings were further strengthened by the observations that the CD103^+^ DC subset from neonatal mice show poor MHC-I presentation of OVA peptides (i.e., SIINFEKL), which was potentially upregulated in the flagellin adjuvanted group in the draining lymph nodes (Fig. 6C). Together, these studies show that TLR5 signaling plays an important role in the maturation, migration and antigen cross presentation of CD103^+^ DC subsets which possibly resonate with recent findings which highlight the adjuvanticity of TLR5 ligands via the mucosal route (33-35).

**Figure 7.**
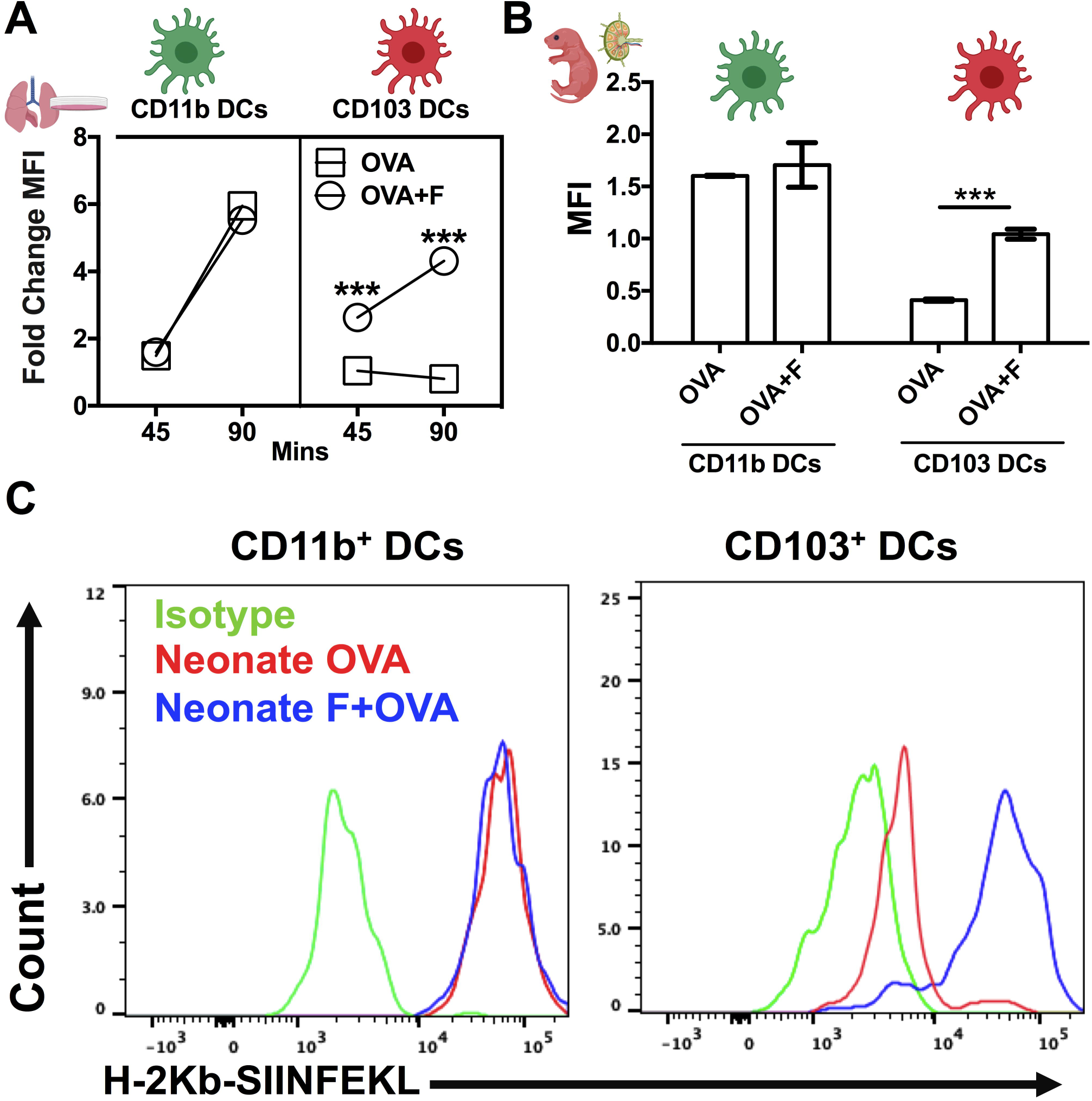
Flagellin regulates phagosome maturation and antigen degradation by neonatal lung CD103^+^ DCs. (**A**) DQ-OVA gives green fluorescence in a green channel, and the co-emergence of a red signal reflects the accumulation of cleaved OVA peptides enabling assessment of cytoplasmic differences in organelle distribution after endocytosis. Stimulation of neonatal lung CD11c^+^ cells in vitro with flagellin measured the phagosome accumulation of DQ-OVA peptides in a time dependent manner. (**B**) Antigen processing capacities are significantly increased by flagellin in isolated neonatal LN migDC subsets, as determined by the MFI ration of OVA^green^ to OVA^red^ signal signal. (**C**) Anti-mouse MHC Class I antibody bound to SIINFEKL specifically reacts with ovalbumin-derived peptide SIINFEKL bound to H-2Kb of MHC class I, but not with unbound H-2Kb or H-2Kb bound with an irrelevant peptide. Draining lymph node CD103^+^ DC subset from the neonates show poor MHC-I presentation of OVA peptides (SIINFEKL), which was reversed in the Flagellin adjuvanted group. n = 5 per age group. ***p < 0.001, ns, not significant, determined by repeated measures two-way ANOVA with Sidak post hoc test).

## Discussion

Respiratory infection is a major cause of mortality in the infants, with on-going studies assessing the underlying mechanisms for the susceptibility of neonates, including immunological differences in the mucosal compartment (2). Neonatal mucosal barriers are bombarded by environmental, nutritional, microbial and pathogenic exposures after birth (36). Study of age-dependent differences of host mucosal immunity and the neonatal immune system may allow us to better understand the establishment of host–microbial homeostasis and responses to pathogens and vaccines. Indeed, the majority of global immunization schedules are focused on the pediatric age group. However, newborns display distinct immune responses, leaving them vulnerable to infections and impairing immunization via the slow initiation, deceased magnitude of immunogenicity, reduced persistence of functional Abs and cell-mediated responses (22). In addition, infectious pathogenesis in neonates might significantly differ from that of older children and adults (22). As vaccine development pipelines do not always rationally tailor formulations (adjuvants, delivery systems etc.) for use in early life, it is important to understand how best used to optimize vaccine efficacy by taking into account early life immune ontogeny (37). For example, early life vaccination against intracellular pathogens has proven difficult (38). Neonates and infants therefore may require different therapeutic vaccine approaches and as compared to the established regimen applied to adults.

Innate immune response in early life, especially at the mucosal site is important not only to provide the protection against infections but also to activate the adaptive arm of immunity and generate the memory cell pool. Neonates typically demonstrate polarized activation of innate immunity with strong Th2 but limited Th1 responses depending on the stimulus, low interaction of migratory DCs with lymphocytes in the lymph nodes, and often-inefficient adaptive responses (31). In the present study, we numerically, phenotypically and functionally characterized the APCs isolated from neonates and adult mice lungs and monitored their response to different PRR stimuli. In the CD11c enriched cells, we identified mature APCs as MHC-II^high^, and four major APC precursors were gated based on the expression of Ly6C^+/−^ and CD11b^+/−^ in the MHC-II^low^ population. Compared to adult mice, neonate mice demonstrated lower frequencies of CD103^+^ and CD11b^+^ migDC subsets, while the percentages of Ly6C^+^ precursor cells were abundant. Low percentages and qualitative differences in migDC subsets correlate with the RSV infection severity in the neonatal mice lungs (31). However, the Ly6C^+^CD11b^+^ monocytic precursor population has not been fully explored for the transcriptional and functional characteristics to identify the molecular pathways, which can be targeted to derive the mature APC and cDC populations. Remarkably, we noted that neonatal mice showed a lower percentage of type-I interferon secreting pDCs, while there were no differences in the antigen uptake capacities of all the APC subsets isolated from neonates and adults. These observations are consistent with previous reports which demonstrated that neonatal lung contains exceptionally low percentage of pDCs while the same cell type is more abundant in the spleen of same mice, and constitute ∼40% of the mature APCs (39). The quantitative differences of the migDC subsets in the lungs were also complemented by the phenotypic deficiencies of their maturation as they displayed the lower expression of maturation markers CD40, CD80 and CD86. Based on these observations, we speculate that the low abundance or immature phenotype of migDCs and pDCs in the lung might be primarily responsible for the higher susceptibility to respiratory infection in early life.

For functional characterization of these APC subsets, we stimulated CD11c enriched cells with different PRR agonists and investigated the expression of maturation markers. Using *in vitro* bone marrow-derived DCs from neonates and adult, we previously identified STING agonist (3,5cGMP) as an adjuvant at an early life (∼ 7 days), which expanded GC B cells and subsequent antigen specific antibody titers, and IgG2c class switching as early as 42 days of life (24). However, the mucosal CD11c^+^ DCs from neonates and adult showed a different pattern of activation to the same stimuli, and flagellin was a most active agonist *in vitro*. While responses to PRR agonists PAM3SK, PAM2CSK, and PolyI:C was impaired, we noted that the MPLA, CL075, CpG and STING induced robust activation of lung DCs from mice lungs. Adjuvant-driven activation of migDC is of interest as these cells not only produce cytokines and induce leukocyte chemotaxis, but also migrate to the draining lymph nodes and initiate the adaptive arm of immunity.

Similarly, among all the stimuli employed, TLR5 agonist flagellin activated neonatal migDCs more robustly and upregulated the expression of lymph node homing receptor CCR7 on neonatal migDCs. These findings corroborate many studies which highlight flagellin as a potent mucosal adjuvant and highlight its role in the cytokine release, leukocyte chemotaxis, cellular interactions, and the generation of protective innate immunity in the mucosal compartment (40,41). To track migDC mobilization to lymph nodes, we administered DQ-OVA intranasally to neonates and adult mice. DQ-OVA not only acts as an antigen, but is also a very useful tool to study the antigen processing and presentation *in vivo*. While unstimulated DCs from neonates showed poor mobilization from the lung to lymph nodes in response to DQ-OVA antigen, and flagellin co-administration strongly potentiated the migration of both the subsets and also upregulated the expression of CD80 and CD86 on CD103^+^ DC in the lymph nodes. These observations further strengthen our findings and highlight flagellin as a potent mucosal adjuvant in our early life intranasal vaccination model.

Another intriguing finding of our study is the differences in the lysosomal processing of antigens at an early age. DQ-OVA proteolytic cleavage emits the green fluorescence in the endosome, while the accumulation of OVA peptides in the perivascular compartment gives a strong red signal. While there were no differences in the antigen uptake across both the subsets, neonatal CD103^+^ DCs showed very rapid cleavage and clearance of DQ-OVA and significantly lower signal in the red channel, which signifies poor antigen cross presentation *in vivo*. We also looked at the antigen processing capacities of isolated lung migDC subsets, and observed a striking difference in the DQ-OVA processing which signify the endosomal defects in the cross-presenting machinery of CD103^+^ DCs. However, flagellin stimulation revived phagosomal maturation and overcame the slow perivascular accumulation of antigens. We also studied OVA peptide presentation on MHC-I using anti-SINFFKEL antibodies specific for ik-ab *in vivo*, and our results show that flagellin co-administration significantly increased the antigen presentation on MHC-I. These findings suggest that TLR5 signaling in CD103^+^ DC plays an important role in the priming of antigen cross-presentation. However, there is also evidence that flagellin-OVA fusion proteins may promote antigen cross-presentation independently of TLR5 or MyD88 signaling, by facilitating antigen processing, which is presumably depend upon the maturation stage of DCs (42).

While dealing with the continual process of immune education during postnatal development, early life lung-resident immune cells are faced with the distinct challenge of reacting to pathogenic and non-pathogenic exposures in a manner that protects the organ’s critical gas exchange machinery. This “neonatal window of opportunity” is such that early life priming can set the stage for life long host-microbial interaction and immune homeostasis (43-46). Additionally, studies are increasing focusing on whether the composition of early life microbiome may ultimately affect vaccine efficacy (47). For example, several theories are emerging regarding immune ontogeny and exposure during infancy in relation to immune perturbations such as pathogen exposure and vaccination. These include a) the “hygiene hypothesis” (48), b) the “appropriate exposure hypothesis” (49), c) the “founder hypothesis” (50), and d) “trained immunity” and “local innate immune memory” in the lung (51,52). Interestingly, human cDC1 and cDC2 subset distribution is a function of tissue site and basal cDC2’s preferentially exhibit maturation and migration phenotypes in mucosal-draining lymph nodes of adults and elders (53). However, this overabundance of cDC2:cDC1 ratio is a) largely confined to intestinal sites in early life, and b) cDC2 maturation dominance only emerges in the lung in later infancy (∼3 months of life) and is maintained throughout life thereafter. Our studies provide further support for an ontologically driven cDC2:cDC1 ratio in the lung, and refine it to specially highlight a cDC1 dominance in the neonatal period that is supplanted by cDC2 subsets in infancy.

Overall, our study features several strengths, including (a) the first immunophenotypic characterization of murine neonatal lung APC subsets; (b) an unbiased screening of PRR agonists for activity toward neonatal lung DCs; and (c) identification of flagellin as an adjuvant active towards neonatal leukocyes *in vitro* and *in vivo* with potential utility in enabling early life intranasal immunization. Our study also has limitations, including (a) the neonatal lung APC model represents a mix of cells isolated by mechanical and enzyme treatment *in vitro* such that they may not fully reflect *in vivo* biology, (b) the potential effects of flagellin on migDC1 to lymph organs are intriguing but are at this stage inferential, (c) although our studies demonstrated cDC1 maturation with flagellin adjuvanted intranasal ova vaccination, future functional studies (e.g., pathogen challenge) are required to assess the efficacy of this adjuvantation system, and (d) due to species specificity, results in mice may not accurately reflect those in humans.

Overall, to our knowledge, our study is the first to functionally characterize mucosal APC subsets by age and evaluate their responses to different PRR agonists which can be used as mucosal adjuvants and/or potentiators of innate immunity enhance resistance to respiratory infections in early life. We also highlight that the neonatal phase of murine development features a distinct endosomal compartment in mucosal CD103^+^ DCs, specifically a lack of adult like phagosomal maturation and cross-presenting machinery, which could be functionally enhanced by TLR5-mediated regulation. Future studies should further address additional mechanistic differences in the endosomal compartment as well as the effect of flagellin adjuvantation on antigen immunogenicity *in vivo*, including to live viral challenge.

## Acknowledgments

We authors thank the members of the BCH *Precision Vaccine Program* for their helpful discussions. The authors are grateful for the mentorship and support of Drs. Michael Wessels, Francesco Borriello, Dennis Kim and Gary R. Fleisher and *Precision Vaccines Program* Coordinator Diana Vo for administrative and programmatic support.

## Disclosures

The other authors declared no conflict of interest.

