## Supplemental Table 1 for "The TLR5 agonist flagellin modifies phenotypical and enhances functional activation of lung mucosal antigen presenting cells in neonatal mice"

| **Table I. Antibodies used for flow cytometry study of APC subsets** | | | | |
| --- | --- | --- | --- | --- |
| **Anti-mouse** **Abs** | **Fluorochrome** | **Clone** | **Cat. No** | **Company** |
| CD11c | PerCp | N418 | 117326 | Biolegend |
| F4/80 | Pacific Blue | BM8 | 123124 | Biolegend |
| CD11b | PE-Cy7 | M1/70 | 101216 | Biolegend |
| CD103 | APC | 2E7 | 121414 | Biolegend |
| PDCA1 | BV510 | 927 | 747607 | BC Biosciences |
| MHCII | eFluor780 | M5/114.15.2 | 107628 | Biolegend |
| Ly6C | PE | HK1.4 | 128008 | Biolegend |
| CD80 | FITC | 16-10A1 | 104706 | Biolegend |
| CD86 | PE | GL-1 | 105008 | Biolegend |
| CD40 | FITC | HM40-3 | 102906 | Biolegend |
